# Deep learning-based auto-segmentation of swallowing and chewing structures

**DOI:** 10.1101/772178

**Authors:** Aditi Iyer, Maria Thor, Rabia Haq, Joseph O. Deasy, Aditya P. Apte

## Abstract

**Purpose:** Delineating the swallowing and chewing structures in Head and Neck (H&N) CT scans is necessary for radiotherapy treatment (RT) planning to reduce the incidence of radiation-induced dysphagia, trismus, and speech dysfunction. Automating this process would decrease the manual input required and yield reproducible segmentations, but generating accurate segmentations is challenging due to the complex morphology of swallowing and chewing structures and limited soft tissue contrast in CT images.

**Methods:** We trained deep learning models using 194 H&N CT scans from our institution to segment the masseters (left and right), medial pterygoids (left and right), larynx, and pharyngeal constrictor muscle using DeepLabV3+ with the resnet-101 backbone. Models were trained in a sequential manner to guide the localization of each structure group based on prior segmentations. Additionally, an ensemble of models was developed using contextual information from three different views (axial, coronal, and sagittal), for robustness to occasional failures of the individual models. Output probability maps were averaged, and voxels were assigned labels corresponding to the class with the highest combined probability.

**Results:** The median dice similarity coefficients (DSC) computed on a hold-out set of 24 CT scans were 0.87±0.02 for the masseters, 0.80±0.03 for the medial pterygoids, 0.81±0.04 for the larynx, and 0.69±0.07for the constrictor muscle. The corresponding 95^th^ percentile Hausdorff distances were 0.32±0.08cm (masseters), 0.42±0.2cm (medial pterygoids), 0.53±0.3cm (larynx), and 0.36±0.15cm (constrictor muscle). Dose-volume histogram (DVH) metrics previously found to correlate with each toxicity were extracted from manual and auto-generated contours and compared between the two sets of contours to assess clinical utility. Differences in DVH metrics were not found to be statistically significant (p>0.05) for any of the structures. Further, inter-observer variability in contouring was studied in 10 CT scans. Automated segmentations were found to agree better with each of the observers as compared to inter-observer agreement, measured in terms of DSC.

**Conclusions:** We developed deep learning-based auto-segmentation models for swallowing and chewing structures in CT. The resulting segmentations can be included in treatment planning to limit complications following RT for H&N cancer. The segmentation models developed in this work are distributed for research use through the open-source platform CERR, accessible at https://github.com/cerr/CERR.

## 1. INTRODUCTION

Delineating organs at risk (OAR) is central in radiotherapy (RT) treatment planning to limit normal tissue complications following RT. Manual delineation is time-consuming, subjective, and prone to errors due to factors such as organ complexity and level of experience [1]. Therefore, there has been a great deal of interest in automating this process to generate accurate and reproducible segmentations in a time-efficient manner.

In head and neck (H&N) cancer treatment, isolating the larynx and pharyngeal constrictor muscle is of particular interest to limit speech dysfunction and dysphagia, respectively, following RT [2] [3]. The masseters and medial pterygoids have also been identified as critical structures in limiting radiation-induced trismus [4]. However, delineating these structures is challenging due to their complex morphology and the low soft tissue contrast in CT images.

Few semi-automatic and automatic methods have been previously developed to segment OARs in the H&N. Conventional multi-atlas-based auto-segmentation (MABAS) methods involve propagating and combining manually-segmented OARs from a curated library of CT scans through image registration as in [5] and [6]. Other approaches offer strategies to further refine MABAS using organ-specific intensity [7] and texture [8] features or shape representation models [9] [10]. However, MABAS is sensitive to inter-subject anatomical variations as well as image artifacts, and image registration is computationally intensive, requiring several minutes even with highly efficient implementations [11].

Convolutional neural networks (CNNs) have recently been applied successfully to various medical image segmentation tasks. Ibragimov et al. [12] trained 13 CNNs, applied in sliding-window fashion to segment H&N OARs including the larynx and pharynx, which were further refined using Markov Random Field (MRF)-based post-processing. In [13], Ward van Rooij et al. employed the popular 3D U-net [14] architecture to segment H&N OARs including the pharyngeal constrictor muscle. Zhu et all. [11] extended the U-Net model by incorporating squeeze-and excitation (SE) residual blocks and a modified loss function to improve segmentation of smaller structures such as the chiasm and optic nerves. In [15], Men et al. segmented nasopharyngeal tumor volumes in H&N CT using a modified version of the VGG-16 [16] architecture, replacing fully-connected layers with fully-convolutional layers and introducing improved decoder networks to rebuild high-resolution feature maps. Tong et al [17] trained a fully convolutional neural net (FCNN), incorporating prior information by training a shape representation model to regularize shape characteristics of 9 H&N OARs. The FocusNet [18] developed by Gao et al. utilizes multiple CNNs to segment H&N OARs including the larynx, first segmenting large structures, then segmenting smaller structures with specifically designed sub-networks.

In this work, we present a fully automatic method to segment swallowing and chewing structures and examine its suitability for clinical use. To the best of our knowledge, this is the first [35] deep learning-based method for segmenting the chewing structures in CT images. We propose a novel framework in which DeepLabV3+ segmentation models are trained sequentially to guide the localization of each structure group based on previously-segmented structures. Model ensembles are created using three orthogonal views (axial, sagittal, and coronal) in 2.5D and shown to improve segmentation accuracy of H&N OARs compared to models trained on a single-view.

## 2. MATERIALS AND METHODS

### 2.1 Dataset

CT scans of 243 H&N cancer patients from our institution were retrospectively collected under IRB 16-142 and accessed under IRB 16-1488 to develop the auto-segmentation methods in this work. The dataset was randomly partitioned into training (80%), validation (10%), and testing (10%) sets. The validation set (10%) was used to estimate model performance while tuning hyperparameters, and the testing set (10%) was used for unbiased evaluation of the final model. Representative CT images showing considerable variation in head pose, shape, and appearance around the structures of interest, including slices with dental artifacts due to dental implants, were included in the dataset. Additional data characteristics are listed in Table 1. Manual segmentations of the masseters (left, right), medial pterygoids (left, right), larynx, and constrictor muscle, generated using MSKCC’s in-house treatment planning system, served as the reference standard. Reference contours of the masseters and medial pterygoids were available in 60% of the scans, and the constrictor muscle in 97% of the scans (larynx was provided in all scans).

**Table 1.**
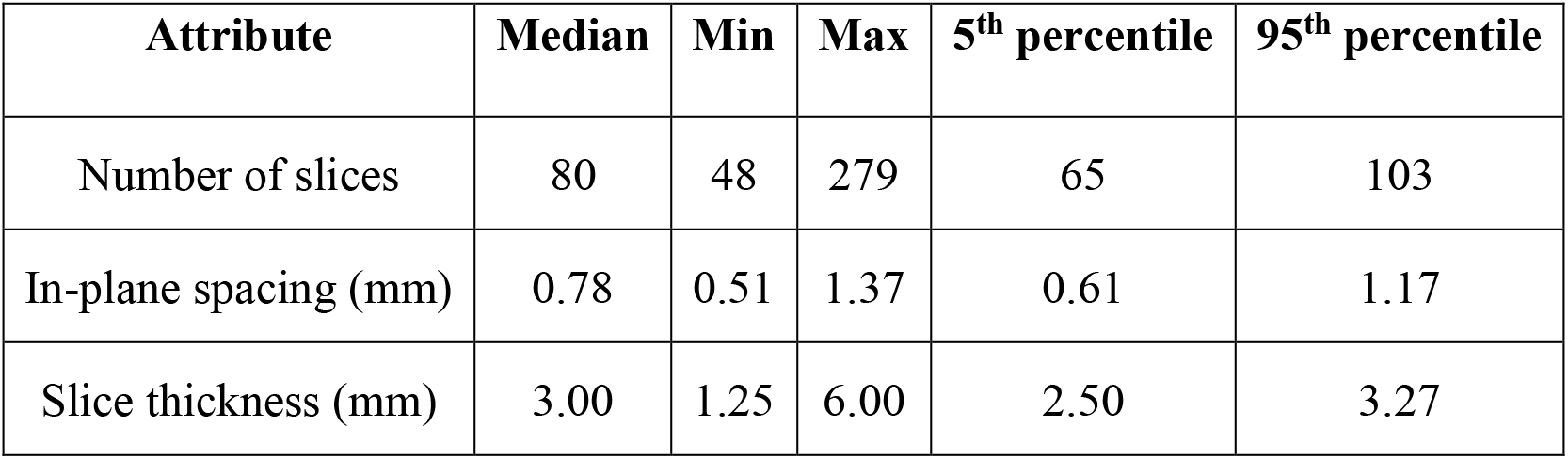
Summary of data characteristics for axial images

| Attribute | Median | Min | Max | 5<sup>th</sup> percentile | 95<sup>th</sup> percentile |
| --- | --- | --- | --- | --- | --- |
| Number of slices | 80 | 48 | 279 | 65 | 103 |
| In-plane spacing (mm) | 0.78 | 0.51 | 1.37 | 0.61 | 1.17 |
| Slice thickness (mm) | 3.00 | 1.25 | 6.00 | 2.50 | 3.27 |

### 2.2 CNN segmentation model

The DeepLabV3+ CNN [19] was selected due to its impressive performance on the PASCAL VOC 2012 and Cityscapes datasets. Moreover, it has previously been applied successfully to medical image segmentation tasks, e.g., by Elguindi et al. [26] to segment the prostate in MR images and by Haq et al. [27] to segment cardio-pulmonary substructures in CT images. This architecture offers the advantage of an encoder network that can capture contextual information at different scales using multiple dilated convolution layers applied in parallel at different rates and a decoder network capable of effectively recovering object boundaries.

Here, training was performed using ResNet-101 [20] as the backbone of the encoder network. A publicly-distributed implementation [21] of DeepLabV3+ using the Pytorch [33] framework was utilized, and a soft-max layer was appended to obtain voxel-wise probabilities. On training a single multi-class model to segment all the structures of interest, it was observed that the imbalance in class labels due to large differences in OAR sizes led to poor performance on smaller structures. Augmenting the loss function based on class frequencies did not produce significant improvement, as previously observed in [18]. Consequently, three separate models were developed-one to segment the chewing structures and one each for the larynx and the constrictor muscle. A sequential segmentation strategy (figure 1) was utilized in which each segmented OAR was used to constrain the location of subsequently segmented OARs. Additionally, an ensemble of three models was developed per OAR group using 2D axial, sagittal, and coronal slices. As compared to 3D convolutional neural networks (CNNs) which are highly memory-intensive, training in 2D enabled us to employ a more complex CNN while still providing contextual information and increased redundancy from the three different orientations.

**Figure 1.**
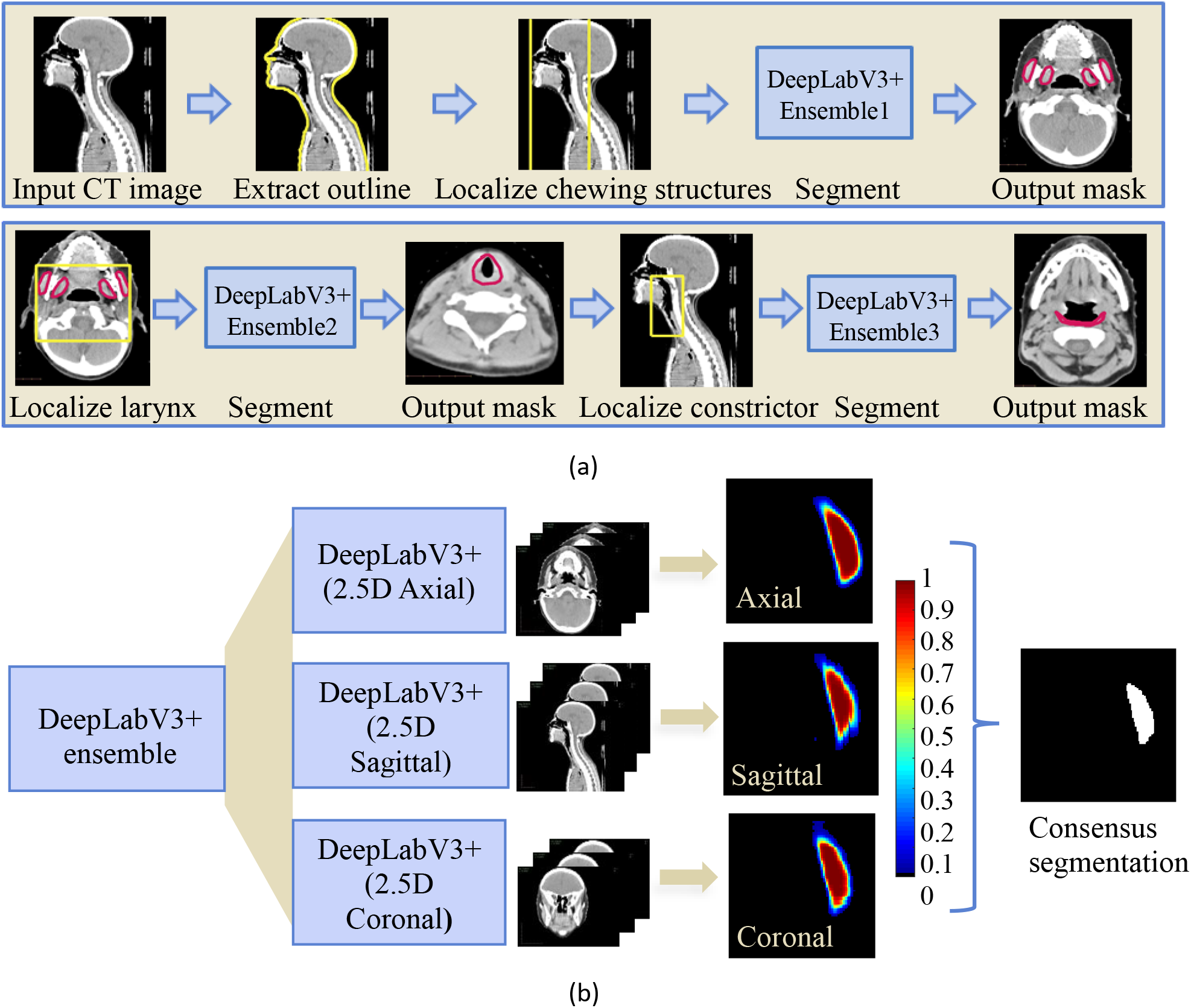
(a) Sequential framework for segmenting chewing and swallowing structures using deep learning, in which each segmented OAR group is used to improve localization of subsequently segmented OARs. (b) Example showing consensus segmentation of left masseter using ensemble model trained on 3 orthogonal views (axial, sagittal, and coronal).

**Figure 2.**
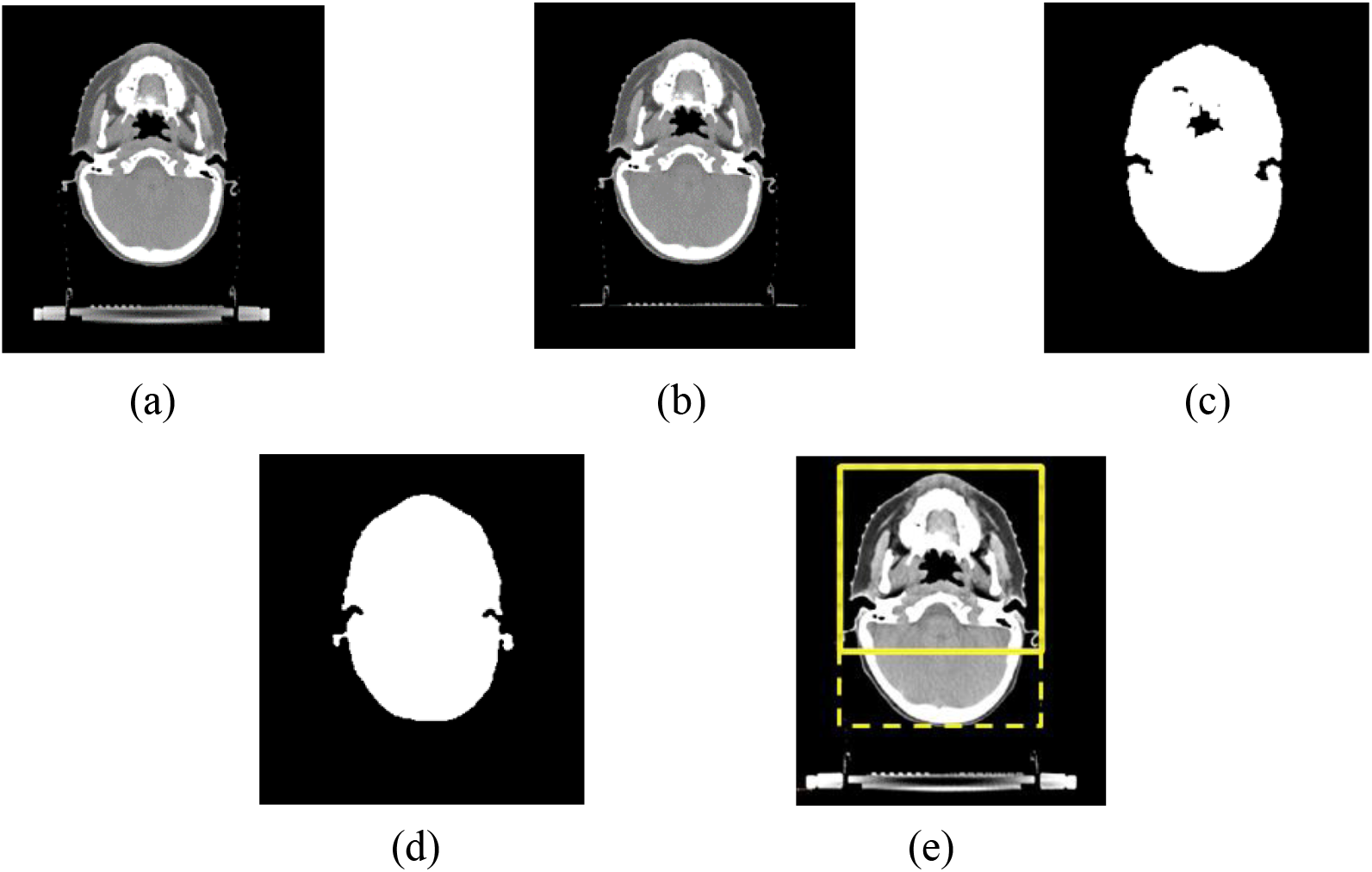
Illustration of method to localize chewing structures in axial H&N CT scans. (a) Input CT image (b) Hough transform-based couch segmentation (c) Intensity thresholding to extract patient outline (d) Morphological post-processing (e) Bounding box with reduced posterior extent.

A high-performance cluster with four NVIDIA GeForce GTX 1080Ti GPUs with 11GB of memory each was used in training. Batch-normalization was applied using mini-batches of 8 images, resized to 320 x 320 voxels. Input images were standardized by scaling the intensities to [0,1] and normalizing to zero-mean and unit-variance. Data augmentation was performed through image scaling, cropping, and rotation. Input channels were populated using three consecutive slices (axial, sagittal, and coronal, respectively, for the three models), and the cross-entropy loss was employed in training. Data preprocessing and export to HDF5 [28] format was performed using the Computational Environment for Radiological Research (CERR) [23]. Details of the individual models are presented in the following sections, and Table 2 summarizes the hyperparameters used.

**Table 2.**
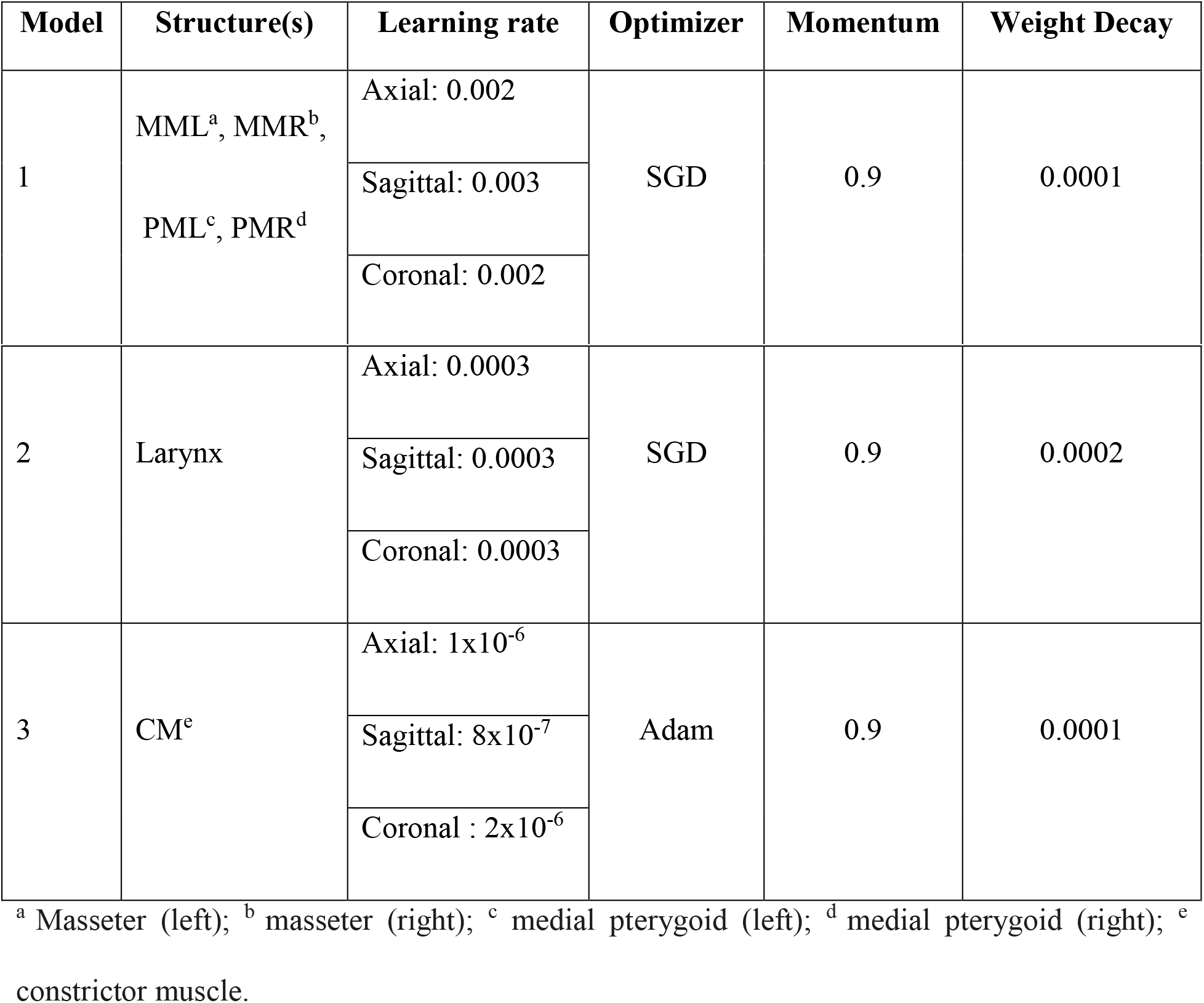
Summary of hyperparameters for training.

### 2.3 Chewing structures

A multi-label model was trained to segment the left and right masseters and medial pterygoids. H&N CT scans were automatically cropped around the patient’s outline prior to training. The couch was first detected using the Hough transform and masked out. This was followed by intensity-based thresholding and morphological post-processing to extract the patient’s outline. A bounding box was generated around the outline and its posterior extent was limited by 25%. 2D axial, sagittal, and coronal images and corresponding masks of the chewing structures were extracted within these extents for training.

Model weights were optimized using Stochastic Gradient Descent (SGD) with hyperparameters listed in Table 2 and learning rate was decayed following the polynomial scheduling policy. An early stopping strategy was employed to avoid overfitting if there was no improvement in the validation loss.

### 2.4 Swallowing structures

#### 2.4.1 Larynx

CT scans were cropped further, with the anterior, left, right, and superior limits defined by the corresponding extents of the previously segmented chewing structures, as shown in figure 3. 2D axial, sagittal and coronal images and corresponding masks of the larynx were extracted within the resulting bounding box and optimization was performed using SGD.

**Figure 3.**
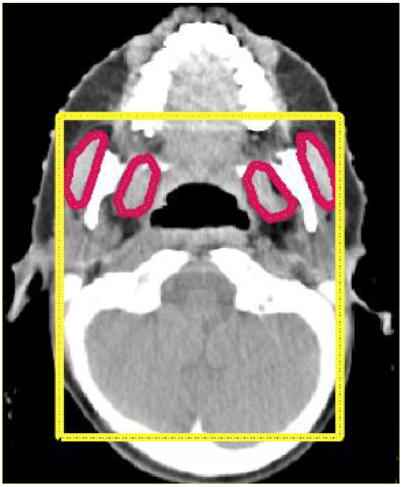
Bounding box (yellow) for localization of larynx using previously segmented chewing structures (red).

#### 2.4.2 Constrictor muscle

To localize the constrictor muscle, CT scans were cropped with the anterior, left, right, and superior limits defined by the corresponding extents of the chewing structures. The posterior and inferior limits were defined by corresponding extents of the larynx, with sufficient padding. 2D axial, sagittal, and coronal images and corresponding masks of the constrictor muscle were extracted within the resulting bounding box, shown in figure 4. Optimization was performed using adaptive moment estimation (Adam) [22].

**Figure 4.**
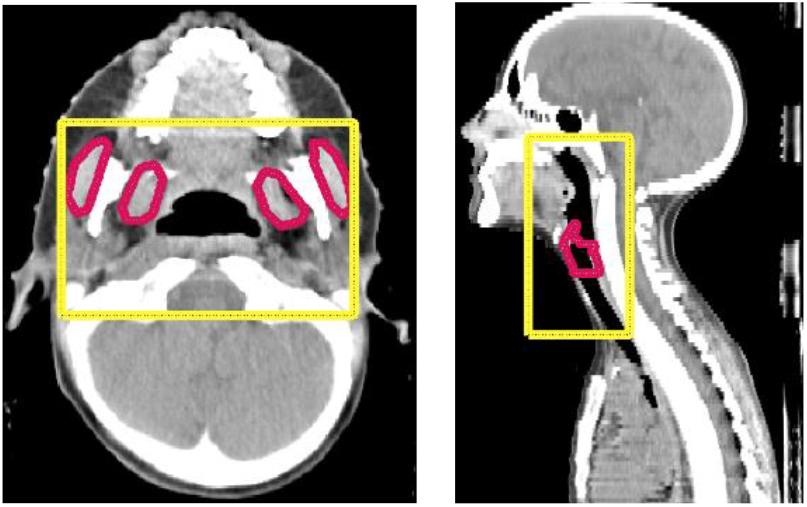
Bounding box (yellow) for localization of constrictor muscle using previously segmented chewing structures and larynx (red).

For each of the above OARs, probability maps returned by the three models (axial, sagittal, and coronal) were averaged and voxels were assigned to the class with the highest resulting probability to produce stable consensus segmentations, robust to occasional failures of the individual models. The segmentation masks were further post-processed to remove isolated voxels by discarding all but the largest connected component. Morphological processing was performed to fill holes and obtain smooth contours.

## 3. RESULTS

### 3.1 Comparison to reference standard

The performance of the proposed method was evaluated on the test set of 24 H&N cancer patients by comparing the results against existing manually delineated segmentations. The Dice Similarity Coefficient (DSC) was used to assess the degree of overlap between manual (A) and automated (B) segmentations, computed as:

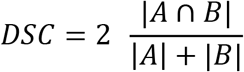

Additionally, the 95^th^ percentile of the Hausdorff distance (HD_95_), i.e., maximum distance between boundary points of A and B, was computed to capture the impact of a few sizeable segmentation errors on the overall segmentation quality. Both DSC and HD_95_ were measured for the individual models (axial, sagittal, coronal) as well as the ensemble to investigate the benefits of a multi-view ensemble.

Examples of segmentations resulting from our algorithm are presented in figure 5 for qualitative assessment. Box plots of DSC and HD_95_ are presented in figure 6.

**Figure 5.**
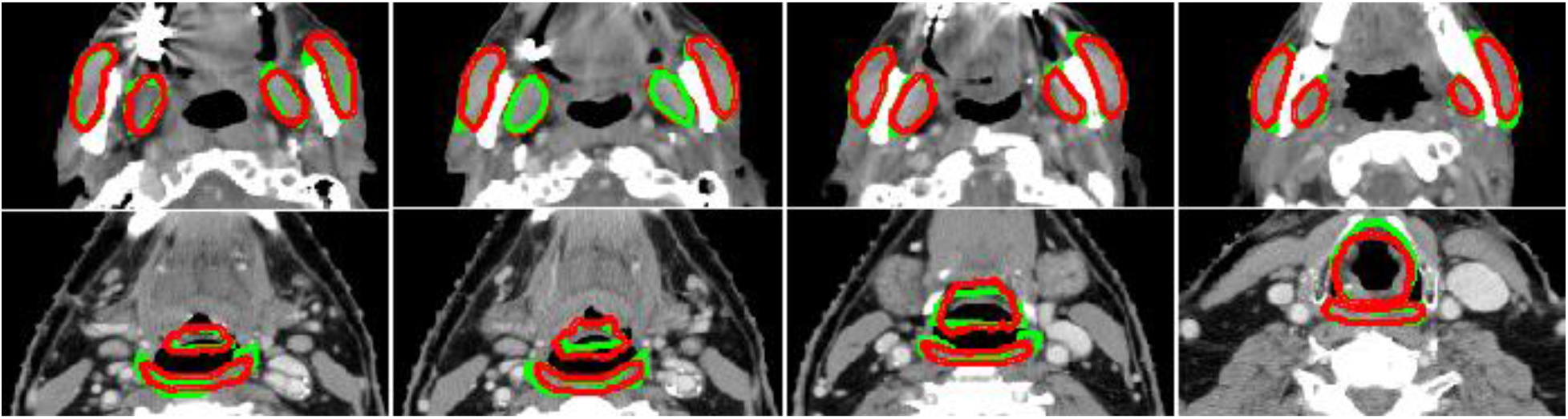
Auto-segmentation results for chewing structures (row-1) and swallowing structures (row-2), shown in four axial cross-sections. Manual reference segmentations are depicted in green and deep-learning-based auto-segmentations in red.

**Figure 6.**
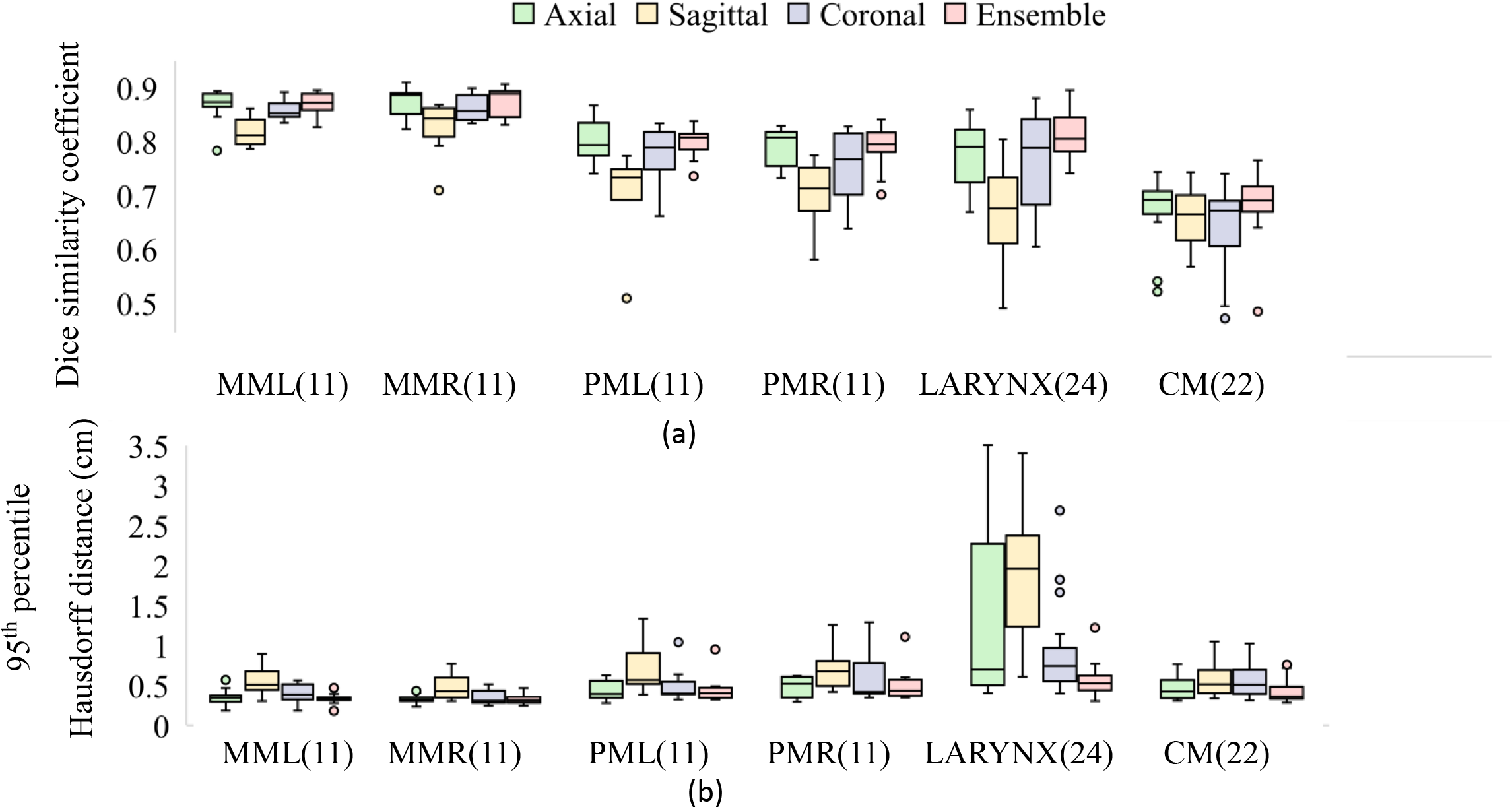
Performance of deep learning models (axial, sagittal, coronal, and ensemble) compared to manual reference segmentations in terms of (a) DSC and (b) HD_95_. Of the 24 test patients, the number with manual contours available for comparison is noted in parentheses. *MML, MMR: masseters (left and right), PML, PMR: medial pterygoids (left and right), CM: constrictor muscle*.

Differences in mean dose, previously identified [29–30] as a possible factor in radiation-induced complications were also computed between deep learning-based and manual contours, and the Wilcoxon signed-rank test was applied to investigate potential statistical disparities (Table 3). Differences in mean dose were not found to be statistically significant for all tested structures at significance level 5%.

**Table 3.** Comparison of mean doses extracted using manual and auto-generated contours. Median, first and third quartiles of percentage differences are presented.

| Structure | Metric | Difference (%) | p-value | No. test patients |
| --- | --- | --- | --- | --- |
| MML<sup>b</sup> U MMR<sup>c</sup> | Ipsilateral<sup>a</sup> mean dose | -0.01 (-1.20, 1.32) | 1.00 | 11 |
| PML<sup>d</sup> U PMR<sup>e</sup> | Ipsilateral<sup>a</sup> mean dose | 0.53 (-0.77, 1.11) | 0.70 | 11 |
| Larynx | Mean dose | 0.38 (-2.02, 6.59) | 0.33 | 24 |
| CM<sup>f</sup> | Mean dose | 0.15 (-0.49, 1.28) | 0.29 | 22 |

<sup>a</sup>Ipsilaterality for the paired structures was decided based upon the side with the highest dose. <sup>b</sup>
Masseter (left); <sup>c</sup> masseter (right); <sup>d</sup> medial pterygoid (left); <sup>e</sup> medial pterygoid (right); <sup>f</sup> constrictor muscle.

### 3.2 Variability of reference standards

The manual reference segmentations came from various sources: the larynx was delineated for treatment planning by multiple observers with variation in level of expertise; the constrictors and chewing structures were each delineated post-treatment by one of two radiation oncology residents for studying dysphagia and trismus, respectively [29] [30]. Regardless of origin, these contours are referred to as observer-1. A randomly selected subset of 10 CT scans from the testing dataset were then re-segmented by a medical physicist (observer-2) to provide a second reference standard. The agreement between the independent observers was measured in terms of DSC and compared to agreement of each observer with the deep learning-based contours (figure 7). Automated segmentations showed better agreement with each observer as compared to the inter-observer agreement.

**Figure 7.**
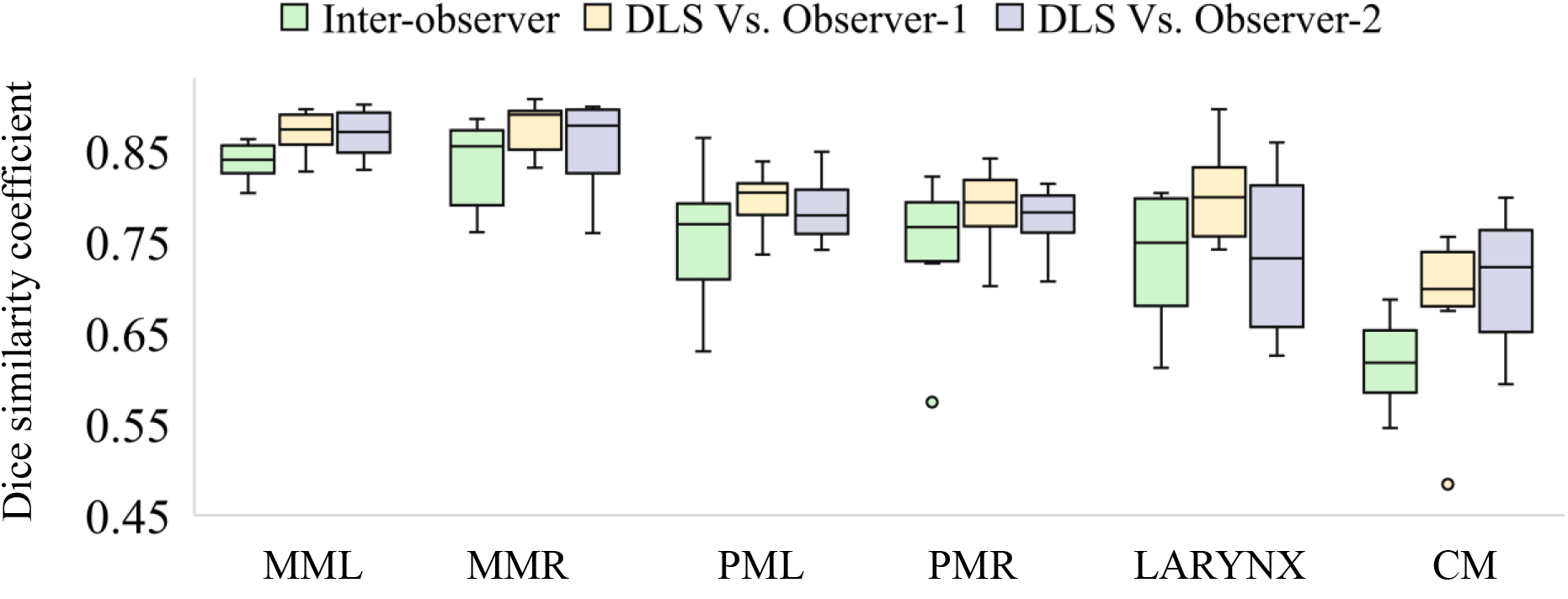
Comparing agreement of deep learning-based segmentations (DLS) with independent observers vs. inter-observer agreement. *MML, MMR: Masseters (left and right), PML, PMR: medial pterygoids (left and right), CM (constrictor muscle)*

### 3.3 Comparison to previously reported methods

We compared the performance of our method with H&N OARs segmentation models from the published literature. These included state-of-the-art CNN models such as the 3D Unet [13], FocusNet [18], a H&N OAR-focused CNN [12], commercial atlas-based segmentation tool SPICE [5] and other atlas-based methods [24,25]. The mean DSCs for the different methods are summarized in Table 4. The performance of our segmentation models matched or exceeded previously presented methods. However, it should be noted that these results were reported on different datasets and do not represent a direct comparison performed on our testing dataset.

**Table 4.** DSC (mean ± std. deviation) for swallowing and chewing structures using the proposed method (column-1) and results from previously published methods.

| Structure | Proposed | van Rooij<br>et al. [13]<br>( 2019 )<sup>1</sup> | Gao et al.<br>[18]<br>( 2019 )<sup>1</sup> | Ibragimov<br>et al. [12]<br>( 2017 )<sup>1</sup> | Tao et al.<br>[5]<br>(2015)<sup>2</sup> | Thomson<br>et al. [24]<br>( 2015 )<sup>2</sup> | Han et<br>al. [25]<br>( 2008 )<sup>2</sup> |
| --- | --- | --- | --- | --- | --- | --- | --- |
| Masseters | 0.87 ± 0.02 | - | - | - | - | - | 0.83 |
| Pterygoids | 0.80 ± 0.03 | - | - | - | - | - | 0.83 |
| Larynx | 0.81 ± 0.04 | 0.78 ± 0.05 | 0.66±0.29 | 0.86 ± 0.04 | 0.73 ± 0.04 | 0.58 | - |
| Constrictor | 0.68 ± 0.07 | 0.68 ± 0.09 | - | 0.69 ± 0.06 | 0.65 ± 0.06 | 0.50 | - |

<sup>1</sup>Deep learning based <sup>2</sup> atlas based.

### 3.4 Distribution of trained models

The segmentation models developed in this work are publicly distributed as a part of CERR’s library of model implementations [31]. Model dependencies were encapsulated using Singularity [34] containers, enabling deployment on a variety of scientific computing architectures and aiding in portability and reproducibility. This also facilitates further analysis through integration with CERR’s radiomics toolbox [32] and dosimetric models [31]. CERR can be downloaded from https://github.com/cerr/CERR. The list of currently available segmentation containers and corresponding links for download are available at https://github.com/cerr/CERR/wiki/Auto-Segmentation-models. It should be noted that the segmentation models are distributed strictly for research use. Clinical or commercial use is prohibited. CERR and containerized model implementations have not been approved by the U.S. Food and Drug Administration (FDA).

## 4. DISCUSSION

We trained three distinct model ensembles to segment structures of different sizes and constrained the location of each structure group based on the extents of previously identified structures. This sequential localization framework was able to handle imbalance in class labels due to differences in the sizes of the OARs. Additionally, we used an ensemble of models trained on three different image orientations to capture important contextual information and provide redundancy in case of failures of the individual models. The performance of the of the multi-view ensemble in terms of DSC either matched or out-performed the individual models and showed lower variance across all structures, as evidenced by tighter bounds in the box plots (figure 6). Further, the multi-view consensus reduced the worst-case segmentation errors (as captured by HD_95_) across all the structures. This suggests that employing a multi-view ensemble may help produce more stable segmentations than single-view models

Of the structures considered, the pharyngeal constrictor muscle was most challenging to segment due to its morphological complexity, high anatomical variability, and low soft tissue contrast. This was reflected in our analysis of manual segmentations by different observers. The lowest inter-observer agreement was observed in delineating the constrictor. In some cases, segmentation was further complicated by tumor infiltration around the periphery. For the larynx, standardizing the reference segmentations (generated by multiple observers with varying levels of experience) prior to training could potentially yield better results. We also observed a few spurious and discontinuous detections of the larynx when using models trained only on axial or sagittal images. This was mitigated by generating a consensus segmentation using information from all 3 views (e.g. figure 8). Finally, the deep learning models for all structures were found to generalize well as the auto-generated results showed good agreement with delineations by a new (unseen) observer.

**Figure 8.**
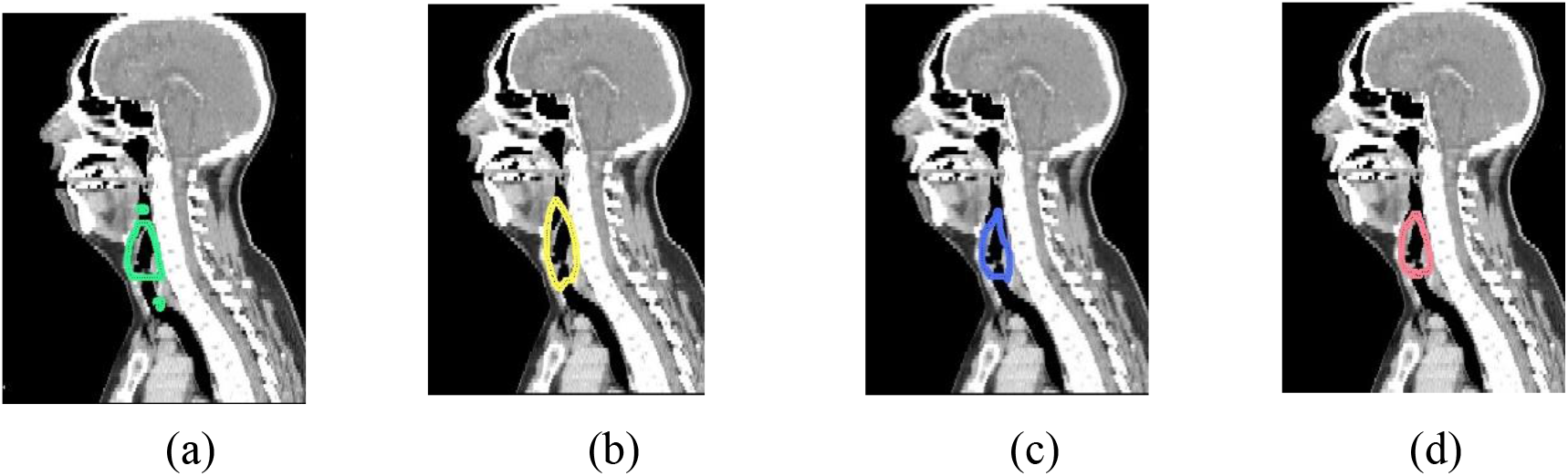
Sagittal cross sections showing auto-segmented larynx using (a) axial only (b) sagittal only (c) coronal only and (d) ensemble models.

We further investigated the potential for clinical application of the trained models by comparing mean doses extracted from manual and automated segmentations. No statistical differences were observed at the 5% significance level.

## 5. CONCLUSIONS

We developed a fully-automatic, accurate, and time-efficient method to segment swallowing and chewing structures in CT images and demonstrated its potential for clinical use. The proposed method introduces a novel framework for sequential localization and segmentation to handle imbalances in OAR sizes. Additionally, the potential advantage of a multi-view ensemble over a single-view model was investigated. As hypothesized, the ensemble models were found to yield more stable segmentations across all structures. This ensemble approach could be applied to improve segmentation quality in other sites as well. The trained models are publicly distributed for research use through the open-source platform CERR using Singularity containers.

## ACKNOWLEDGEMENTS

This work was partially funded by NIH grant 1R01CA198121 and NIH/NCI Cancer Center Support grant P30 CA008748.

